# Structural basis for endotoxin neutralization and anti-inflammatory activity of thrombin-derived C-terminal peptides

**DOI:** 10.1101/232876

**Authors:** Rathi Saravanan, Daniel A Holdbrook, Jitka Petrlova, Shalini Singh, Nils A Berglund, Yeu Khai Choong, Peter J Bond, Martin Malmsten, Artur Schmidtchen

## Abstract

Thrombin-derived C-terminal peptides (TCP) of about 2 kDa are present in wounds, where they exert anti-endotoxic functions. In an effort to elucidate the structural and molecular aspects of these functions, we here employ a combination of nuclear magnetic resonance spectroscopy (NMR), ellipsometry, fluorescence spectroscopy, circular dichroism (CD) measurements, and *in silico* multiscale modeling to define interactions and the bound conformation of a TCP generated by neutrophil elastase, HVF18 (HVFRLKKWIQKVIDQFGE) in complex with bacterial lipopolysaccharide (LPS). In contrast to the disordered state of HVF18 in aqueous solution, its binding to LPS leads to a structural transition, wherein the N-terminus of the peptide forms a unique ß-turn whilst the C-terminus becomes helical. *In silico* modelling and simulations demonstrated that HVF18, as well as related peptides, target the LPS-binding site of CD14, and this interaction was experimentally supported using microscale thermophoresis. Collectively, the results demonstrate the role of structural transitions in LPS complex formation as well as in CD 14 interaction, and provide a molecular explanation for the previously observed therapeutic effects of TCPs in experimental models of bacterial sepsis and endotoxin shock.

**Significance:** Thrombin-derived C-terminal peptides (TCPs) of various sizes are present in human wounds, where they bind bacteria as well as “free” lipopolysaccharide (LPS), and thereby reduce inflammation. In this work, employing a combination of cellular, biophysical and structural studies, combined with *in silico* multiscale modeling, we present the molecular structure of a TCP in association with LPS, and define a previously undisclosed interaction between TCPs and CD14. Further, we show that TCPs exhibit relatively weak but specific affinities, all in the μM range, to both LPS and CD14. These novel structural insights into the function of this class of host-defense molecules will facilitate rational design of novel “dual function” anti-infectives, which target both bacteria and inflammatory signaling.

## Introduction

Lipopolysaccharide (LPS) sensing by Toll-like receptor 4 (TLR4) is crucial in early responses to infection, where an uncontrolled LPS-response gives rise to excessive localized inflammation, such as that found in infected wounds, but also in severe systemic responses to infection [1]. Therefore, although sensing of LPS is important for initial host defense responses, clearance and control of this molecule is critical in order to avoid excessive inflammation and organ damage. For example, LPS triggers NF-κB-mediated up-regulation of tissue factor, and thus formation of thrombin, which in turn activates coagulation and fibrin formation, thereby aiding in initial hemostasis and defense against bacterial invasion [2]. Furthermore, proteolysis of thrombin by neutrophil elastase leads to formation of aggregation prone 11-kDa C-terminal fragments which are present in wounds, and bind to and form amorphous amyloid-like aggregates with both LPS and Gram-negative bacteria, aiding in the subsequent clearance of these aggregates by phagocytosis [3]. In addition, thrombin-derived C-terminal peptides (TCP) of roughly 2 kDa, such as FYTHVFRLKKWIQKVIDQFGE and HVFRLKKWIQKVIDQFGE [4-6], are present in wound fluids, and have been demonstrated to exert anti-endotoxic functions *in vitro* and *in vivo* [4, 7]. Such smaller peptides belong to the diverse family of host-defense peptides (HDP) [8], which includes neutrophil-derived α-defensins and the cathelicidin LL-37 [9, 10], all known to exhibit immunomodulatory activities [11]. Today, there is an urgent need for novel anti-infective therapies [12, 13]. However, current treatments based on antibiotics target bacteria only, and not the accompanying over-activation of immune responses. This uncontrolled stimulation may cause an overwhelming production of inflammatory cytokines leading to systemic inflammation, intravascular coagulation and organ dysfunction, such as seen in sepsis [14, 15]. This is a leading cause of death in the U.S alone, with over 700,000 cases estimated every year, and with mortality rates from 28 to 60% [16, 17]. Treatment concepts based on Nature’s own innate defense strategies, aiming at not only targeting bacteria, but also the excessive immune response, could therefore have a significant therapeutic potential. For instance, many naturally occurring HDPs, as well as fragments of LPS-recognizing proteins, display a strong affinity for LPS, blocking downstream interaction with LPS-binding protein (LBP), thereby reducing cytokine production *in vitro* and *in vivo* [18-20].

Neutralization of circulating LPS by HDPs has been shown to reduce adverse LPS-induced pro-inflammatory effects in experimental animal models [4, 21, 22]. In this context, a prototypic TCP, GKY25 (GKYGFYTHVFRLKKWIQKVIDQFGE), encompassing sequences of natural TCPs previously identified in human wounds [23], has been shown to protect against *P. aeruginosa* sepsis and LPS-mediated shock, mainly via reduction of systemic cytokine responses *in vivo* [4, 24]. These observations, however, disclose neither the exact mode of action, nor whether these peptides can target other molecules apart from LPS. With this background, we set out to characterize the structural prerequisites at the molecular level which underlie the anti-endotoxic actions of such TCPs.

## Results

### Structural considerations and background on peptide structures

The C-terminal region of thrombin of which the sequence GKYGFYTHVFRLKKWIQKVIDQFGE is highlighted in the crystal structure (shown in Fig. 1A), comprises a flexible β-strand segment and a compact amphipathic helix. Previous studies have shown that various proteases can generate TCPs derived from this region [23]. For illustrative purposes, and to highlight the complete, albeit gradual, formation of such fragments, the generation of the 2 kDa TCP HVF18 HVFRLKKWIQKVIDQFGE, cleaved out by human neutrophil elastase [7], is depicted in Figure 1B. A similar generation of the peptide FYT21 (FYTHVFRLKKWIQKVIDQFGE) has been observed after digestion with the bacterial M4 peptidases *Pseudomonas aeruginosa* elastase [6, 23] and *Staphylococcus aureus* aureolysin [23]. Incorporating these endogenous sequences, the prototypic peptide GKYGFYTHVFRLKKWIQKVIDQFGE has been further used in *in vitro* studies on its mode-of-action [24, 25], as well as in several therapeutic *in vivo* studies, aimed at targeting endotoxin-mediated inflammatory responses [4, 7]. The truncated peptide variant VFR12 (VFRLKKWIQKVI) has been shown to constitute a minimal LPS binding site, while not exhibiting any anti-endotoxic effects in cell models [26]. Some key physicochemical parameters of these peptides are indicated in Figure 1C.

**FIGURE 1.**
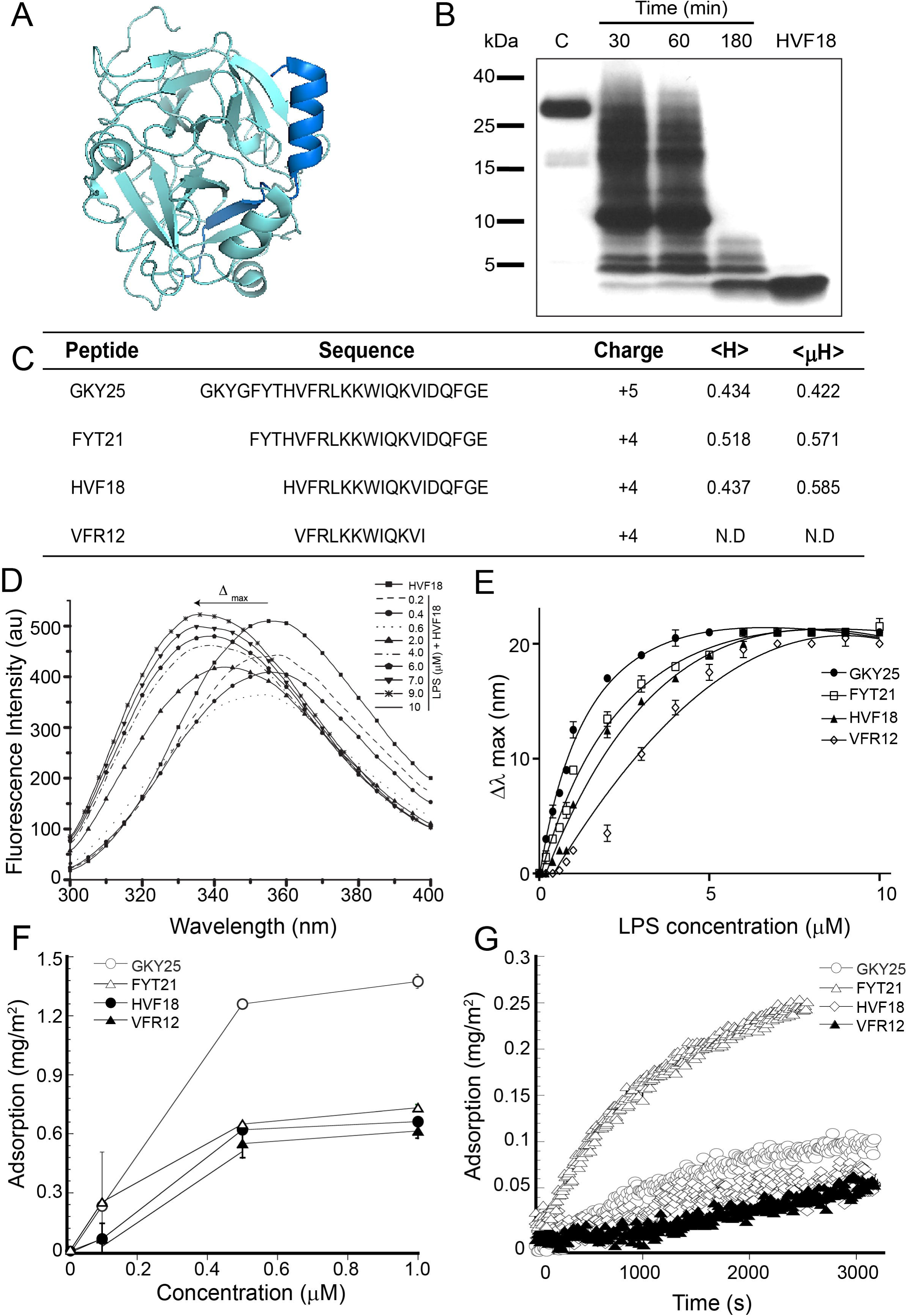
Thrombin C-terminal peptides, and their binding to *E. coli* LPS. (A) 3D model of human α-thrombin highlighting thrombin C-terminal peptide fragment. (B) Western blot analysis of human α-thrombin incubated with neutrophil elastase (NE) for the indicated time period generates thrombin C-terminal peptide fragments of varying lengths. The synthetic peptide HVF18 is loaded as molecular size control corresponding to generated fragments. (C) Table showing thrombin C-terminal peptides amino acid sequences and physicochemical properties determined using Heliquest online server tool. The hydrophobicity and amphipathicity of VFR12 was too short for accurate determination using Heliquest. (D) Intrinsic tryptophan fluorescence emission spectra of HVF18 titrated with increasing concentration of *E. coli* LPS. (E) LPS binding affinity of peptides determined from changes in emission maxima (λ_max_) upon titration, fitted with Graph Pad Prism v7.02 one-site binding model. (F) Ellipsometry showing peptide binding to *E. coli* LPS preabsorbed on solid supports and (G) corresponding peptide adsorption kinetics. All experiments were performed in 10 mM Tris pH 7.4.

### Interactions of TCPs with LPS

Intrinsic tryptophan fluorescence was employed to monitor the interaction of the TCPs with *E. coli* LPS. Figure 1D shows changes in the fluorescence emission spectra of HVF18 as a function of increasing concentration of LPS. In the absence of LPS, the tryptophan residue has an emission maximum at 357 nm. Addition of LPS results in a concentration-dependent blue shift in the emission maximum, indicating peptide binding to LPS. A similar shift in emission maximum was observed for the other TCPs, albeit differences in the extent of the blue shift were noted (Fig. S1). The change in fluorescence emission maxima (λ_max_) of the TCPs was utilized to obtain an indication of peptide binding affinities to LPS (Fig. 1E). As shown in Table 1, GKY25 displayed the highest affinity (*K_d_* =2 ± 0.29 μM) among the peptides investigated, followed by the endogenous fragments HVF18 (*K_d_* = 4.24 ± 0.90 μM), and the shorter VFR12 (*K_d_* =11.45 ± 0.45 μM). Furthermore, the localization of the tryptophan residue in the LPS complexes was determined by quenching of the tryptophan fluorescence with a neutral quencher, acrylamide. Table 1 shows the obtained Stern-Volmer quenching constants (*K_SV_*) of the peptide in unbound and LPS-bound states. Although many factors determine the photophysics of tryptophan, a large *K_SV_* for the peptides in their unbound state generally indicates complete exposure of the tryptophan residue to buffer, while a low *K_SV_* in the LPS-bound states implicates that exposure of the tryptophan to the solvent is restricted owing to incorporation of the tryptophan side chain into the hydrophobic milieu of LPS.

**Table 1.** Binding affinities (*K_d_)* and Stern-Volmer’s quenching constants (K_*SV*_) determined from the intrinsic tryptophan fluorescence measurements.

| Peptide | Kd (μM) | $K_{SV}$ ($M^{-1}$) | |
| --- | --- | --- | --- |
|  |  | Buffer | LPS |
| GKY25 | $2.0 \pm 0.29$ | 29.7 | 4.5 |
| FYT21 | $3.69 \pm 0.70$ | 27.0 | 5.3 |
| HVF18 | $4.24 \pm 0.90$ | 41.8 | 6.6 |
| VFR12 | $11.5 \pm 0.90$ | 35.1 | 6.8 |

By analogy, ellipsometry results indicate that these TCPs bind substantially to LPS (Fig. 1F-G). GKY25 adsorbed to LPS with the highest affinity among the peptides investigated, followed by FYT21, HVF18, and VFR12. At low peptide concentrations, the binding of the TCPs is similar, showing that the initial LPS recognition and adsorption is determined by peptide charge and electrostatic interactions. At increasing peptide concentrations (P:L ratio), peptide length variations affect LPS binding, indicating the importance of hydrophobic interactions required for LPS penetration (Fig. 1F). Taken together, the ellipsometry and tryptophan fluorescence results indicate that TCPs exhibit a peptide length-dependent binding to LPS.

### Effects of TCPs on LPS scavenging and blocking of pro-inflammatory responses

Next, we assessed the ability of the peptides to neutralize free LPS in solution using the highly sensitive limulus amebocyte lysate assay (LAL) assay [27]. As can be seen in Figure 2A, GKY25, FYT21, and HVF18, but not VFR12, neutralized LPS in the LAL assay in the dose range studied, and a length-dependent inhibitory activity was observed, corresponding to the ellipsometry and tryptophan fluorescence data outlined above. Likewise, using LPS-stimulated THP-1 monocytes followed by evaluation of NF-κB activation, a similar blocking of endotoxin responses was observed (Fig. 2B), in which VFR12 did not block NF-κB activation (Fig. S1D). Notably, the peptide FYT21, previously identified in infected wounds [6, 23], blocked LPS-induced pro-inflammatory response and displayed LPS neutralization to a similar extent as GKY25, previously used in therapeutic studies (Fig. 2B). Also notable is that the fragments FYT21 and HVF18 share relevant physiological concentrations for both thrombin and other TCPs [3]. Overall, these data demonstrate a correlation between TCP length, LPS-neutralization, and blocking of pro-inflammatory responses *in vitro*.

**FIGURE 2.**
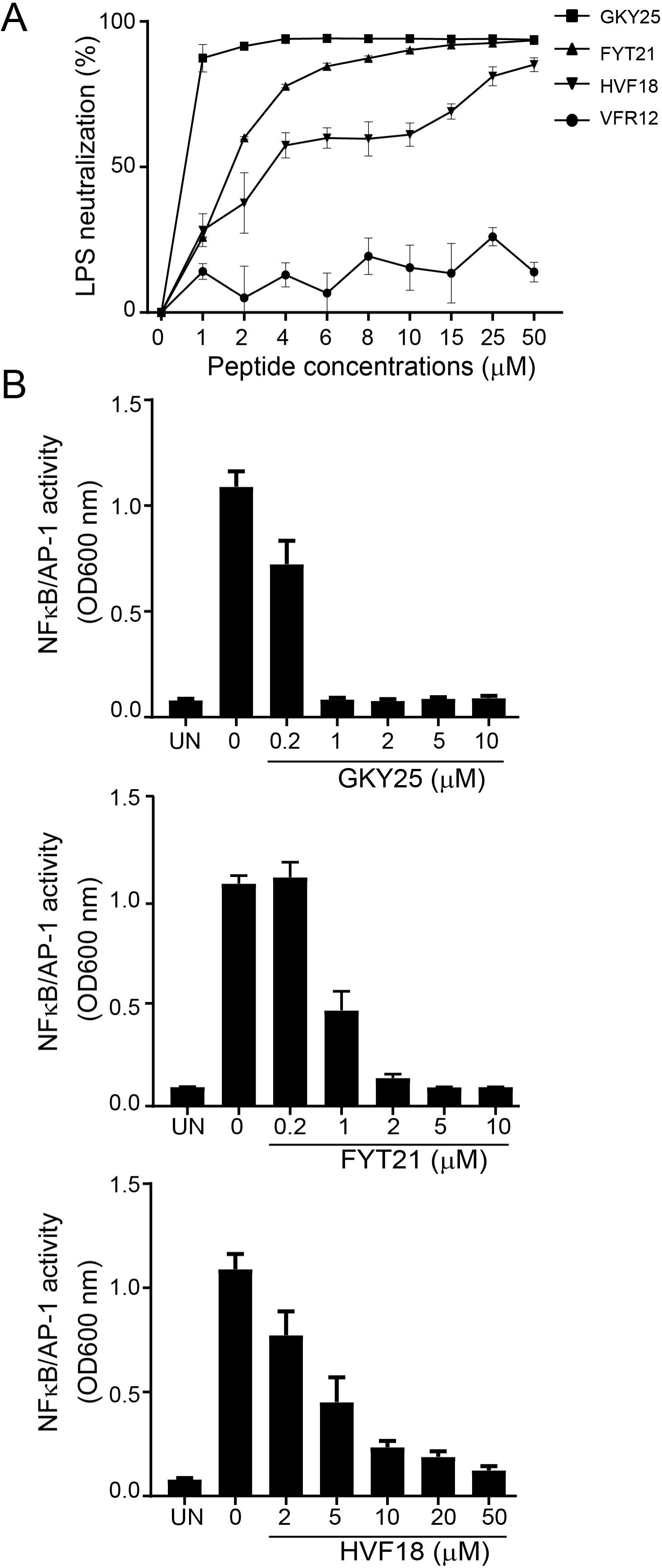
Thrombin C-terminal peptides bind and inhibit LPS-induced proinflammatory responses. (A) Limulus amebocyte (LAL) chromogenic assay showing variations in TCPs ability to neutralize *E. coli* LPS in solution. (B) Inhibition of NF-kB activation in THP-1 monocytes stimulated with 10 ng/ml *E. coli* LPS upon treatment with indicated TCPs at the specified concentrations, 20 hours post stimulation.

### NMR studies of TCPs in free and LPS bound states

NMR structural studies were employed to gain further insight into the molecular interactions of TCP-LPS complex formation. The spin systems in the TOCSY spectra of VFR12 and HVF18 are well dispersed and permit complete assignment of proton resonances (Fig. S2A-B). However, in the case of FYT21 and GKY25, the N-terminal (GKYGFYTHV) proton resonances overlapped or were absent, rendering assignment incomplete (Fig. S2C-D). Changes in D2O (5%) and pH did not improve the spin system assignments. Nonetheless, the NOESY spectra of the free peptides were largely characterized by weak intra-residue (CαH_*i*_/NH_*i*_) and sequential (CαH_*i*_/NH_*i*+1_) connectivities between backbone proton resonances (Fig. S2). The absence of medium-(*i* to *i*+2/ *i*+3/ *i*+4) and long-range nuclear Overhauser enhancements (NOEs), particularly involving aromatic side-chain proton resonances (6.8 to 7.5 ppm) clearly indicate that the TCPs are highly dynamic in solution, consistent with the random coil observation from circular dichroism (CD) spectra (Fig. S2E).

However, a series of 1D ^1^H-^1^H spectra of GKY25 or FYT21 with successive addition of LPS showed prominent chemical shifts in proton resonances of the backbone and side-chain regions (Fig. S3A and S3B). Moreover, visible aggregates formed, indicating that the peptides do not undergo fast-to-intermediate exchanges between free and LPS-bound state within the NMR time-frame, thus precluding three-dimensional structure determination. Nonetheless, in the case of HVF18 or VFR12, addition of LPS yielded concentration-dependent line broadening of the backbone and side-chain proton resonances without any significant chemical shift changes, enabling LPS-bound structure determination using tr-NOESY (Fig. S3C-D).

The tr-NOESY spectra of HVF18 showed a drastic improvement in the number and intensity of NOE cross-peaks, involving backbone and side-chain proton resonances in the presence of LPS (Fig. 3). The down-field shift of the indole proton of W8 (~ 10.17 ppm), as well as of the degenerate aromatic ring protons of Phe (7.35-7.25 ppm), showed several NOE connectivities with aliphatic side-chain proton resonances of residues L5, I9, V12, I13 (Fig. 3A and 3D). Interestingly, NOEs involving the indole proton of W8 with the CαH and CβH of R4, K6, and K7 were also identified. Together, these results demonstrate conformational stabilization and the occurrence of side-chain interactions in LPS-bound HVF18.

**FIGURE 3.**
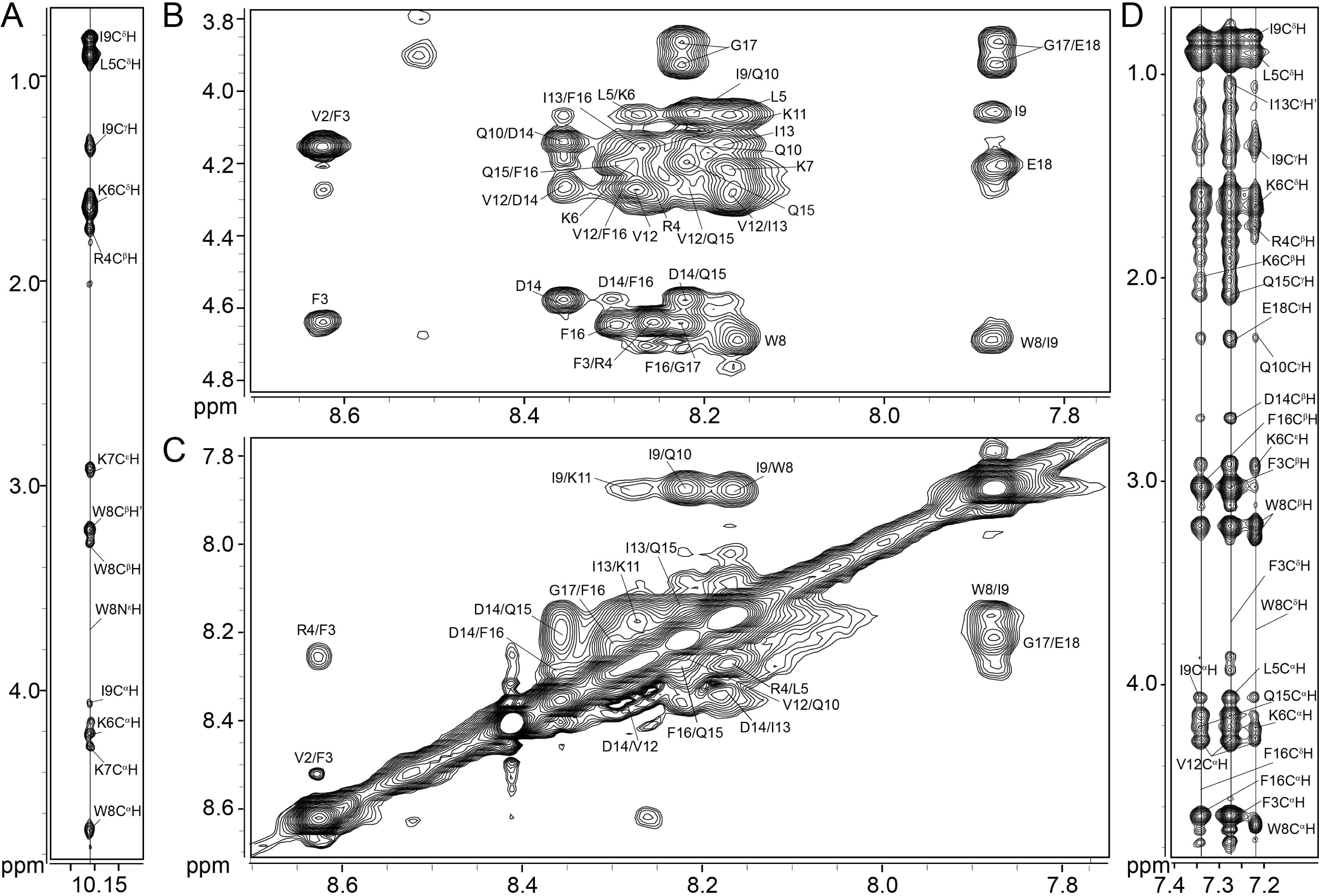
tr-NOESY spectra of HVF18 in LPS micelles. Two dimensional ^1^H-^1^H tr-NOESY spectra of HVF18 (0.5 mM) in the presence of LPS (20 μM) (A) showing NOE connectivities involving W8 indole proton with the upfield shifted aliphatic proton resonances. (B) Finger print region showing sequential and medium range NOE connectivities between backbone alpha and amide proton (CαH/HN) resonances, (C) section showing NOE connectivities among backbone amide (HN/HN) resonances and (D) section showing NOE connectivities between the aromatic ring protons of F3, F16 and W8 with the upfield shifted aliphatic proton resonances. Tr-NOESY spectrum of HVF18 in LPS micelles was obtained in aqueous solution (pH 4.5) at 298K in a Bruker DRX 600 MHz NMR instrument.

Figure 4A summarizes the tr-NOEs of HVF18 observed in the presence of LPS. Intense sequential (CαH_*i*_/NH_*i*+*1*_) as well as backbone NH/NH, NOEs are seen for most residues, except for overlap of the CαH proton resonances with backbone amide proton resonances (ω2) for a few residues. Several medium-range NH/NH (*i* to *i*+2) NOEs and Cα/NH (*i* to *i*+2, *i*+3 and *i*+4) NOEs could be observed for the C-terminal region I9-G17 residues, suggesting an α-helical conformation. In contrast, the N-terminal region H1-W8 is characterized by the presence of strong sequential (CαH_*i*_/NH_*i*+*1*_) NH/NH and a few NH/NH (*i*, *i*+2) NOEs, suggesting a lack of helical conformation at the amino terminus.

**FIGURE 4.**
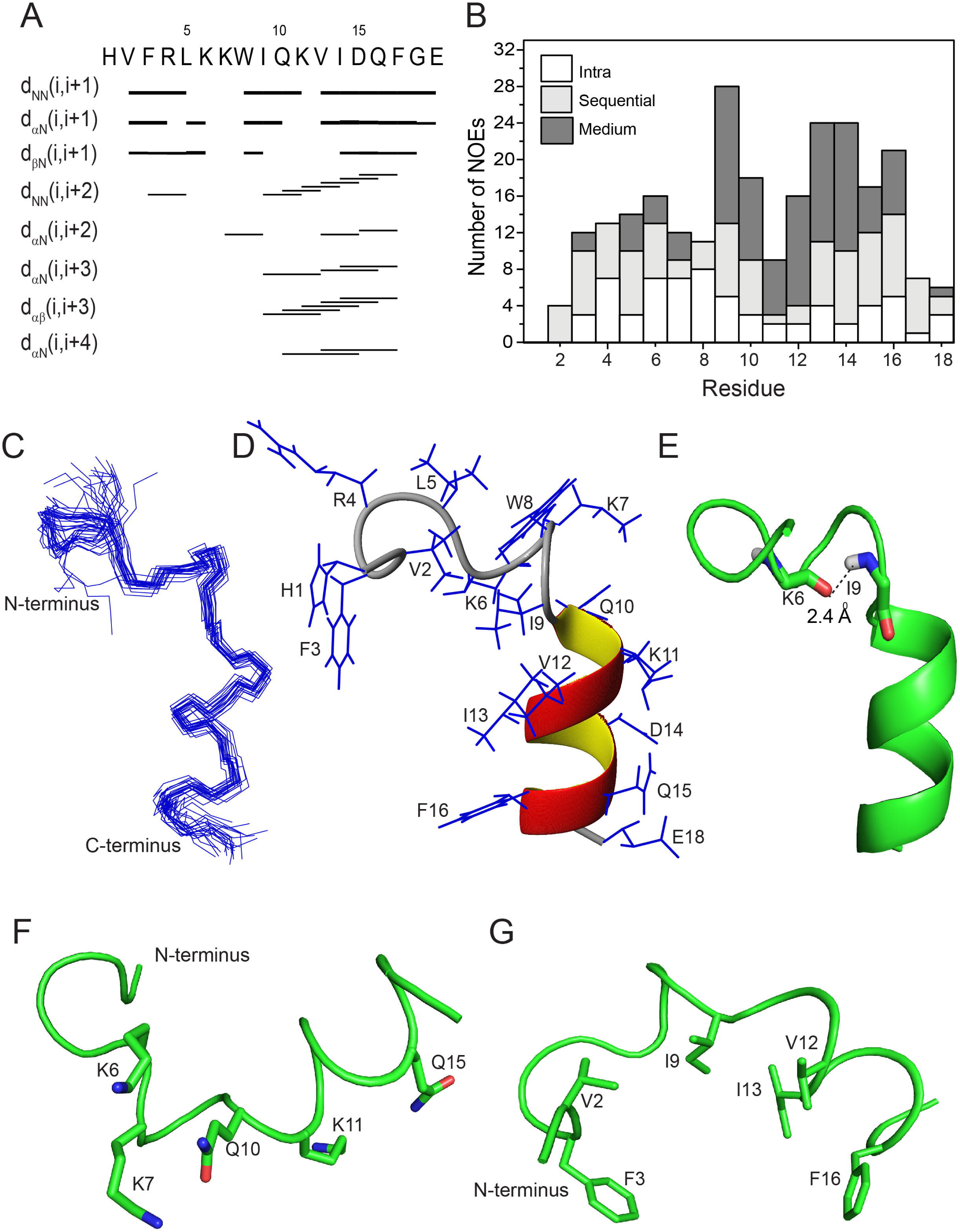
Analyses of tr-NOESY spectra and three dimensional solution structure of HVF18 in LPS micelles. (A) Diagram summarizing the sequential and medium range NOE contacts identified in HVF18. (B) Bar diagram showing number and type of NOEs observed for HVF18 in complex with LPS as a function of amino acids. (C) Superposition of backbone atoms (N, Cα, C’) for the 20 lowest energy conformers of LPS bound HVF18. (D) A representative structure of LPS bound HVF18 showing backbone topology and side-chain dispositions and (E) plausible hydrogen bond formation between K6 and I9 in the β-turn region. Ribbon representation of peptide backbone showing amphipathic disposition (F) of hydrophilic residues along the convex side (G) and the hydrophobic residues on the concave side of a curved backbone fold of HVF18 in complex with LPS. Figures were generated using MolMol v2K.2 and Pymol v1.7.4.0

Likewise, for VFR12 a large number of intense NOE cross-peaks involving backbone and side-chain proton resonances were observed in the presence of LPS (Fig. S4). As above, the continuity of sequential (*i* to *i*+1) NH/NH and Cα/NH NOEs was disturbed due to overlap. However, few medium-range (*i* to *i*+2) NH/NH NOEs and (*i* to *i*+2, *i*+3 and *i*+4) Cα/NH NOEs were observed (Fig. S4B-C). Notably, long-range NOE contacts between the F2 aromatic ring protons with CαH and CβH of W7, as well as NOE contacts between the W7 NεH proton and F2 backbone and side-chain protons, were observed (Fig. S4D). In addition, the indole proton of W7 showed NOE contacts with the aliphatic side-chain proton resonances of residues I8, V11, I12, suggesting close interactions of amino acid side-chains in LPS-bound VFR12. In summary, the analyses of tr-NOESY spectra of HVF18 and VFR12 demonstrate that LPS binding induces backbone stabilization and (partial) formation of α-helical conformation.

### Structure of HVF18 and VFR12 bound to LPS micelles

The 3D structure of LPS-bound HVF18 was determined from a total of 145 tr-NOE-derived distance restraints and dihedral angle restraints (Fig. 4B and Table 2). Figure 4C superposes the backbone atoms of the 20 lowest energy structures of HVF18. The structure is well-defined, with a backbone and heavy atom RMSD of 0.88 ± 0.41 and 1.73 ± 0.46 Å, respectively. In the LPS-bound state, the C-terminus of HVF18, encompassing residues Q10-G17, forms a compact helix. In contrast, the N-terminal residues, F3-L5 are in an extended state, followed by a β-turn involving residues K6-K7-W8-I9 (Fig. 4D). The presence of strong backbone HN/HN (i to i+1) NOEs, medium intensity Cα/NH (i to i+2), alongside the absence of Cα/NH (i to i+3) NOEs, confirms the formation of type II β-turn (Fig. 4A). Furthermore, the close proximity of the K6 carbonyl oxygen and the amide proton of I9 (2.2 -2.4 Å), in all calculated structures indicate the occurrence of hydrogen-bond formation (Fig. 4E). The β-turn gives the peptide backbone a curved shape with amphipathic side-chain dispositions. The side-chains of R4/K6/K7 and Q10/K11/Q15 form two hydrophilic clusters connected by the central hydrophobic W8 residue (Fig. 4F) on the convex side, whereas the side-chains of V2/F3/I9/V12/I13/F16 form a concave hydrophobic core (Fig. 4G).

**Table 2.**
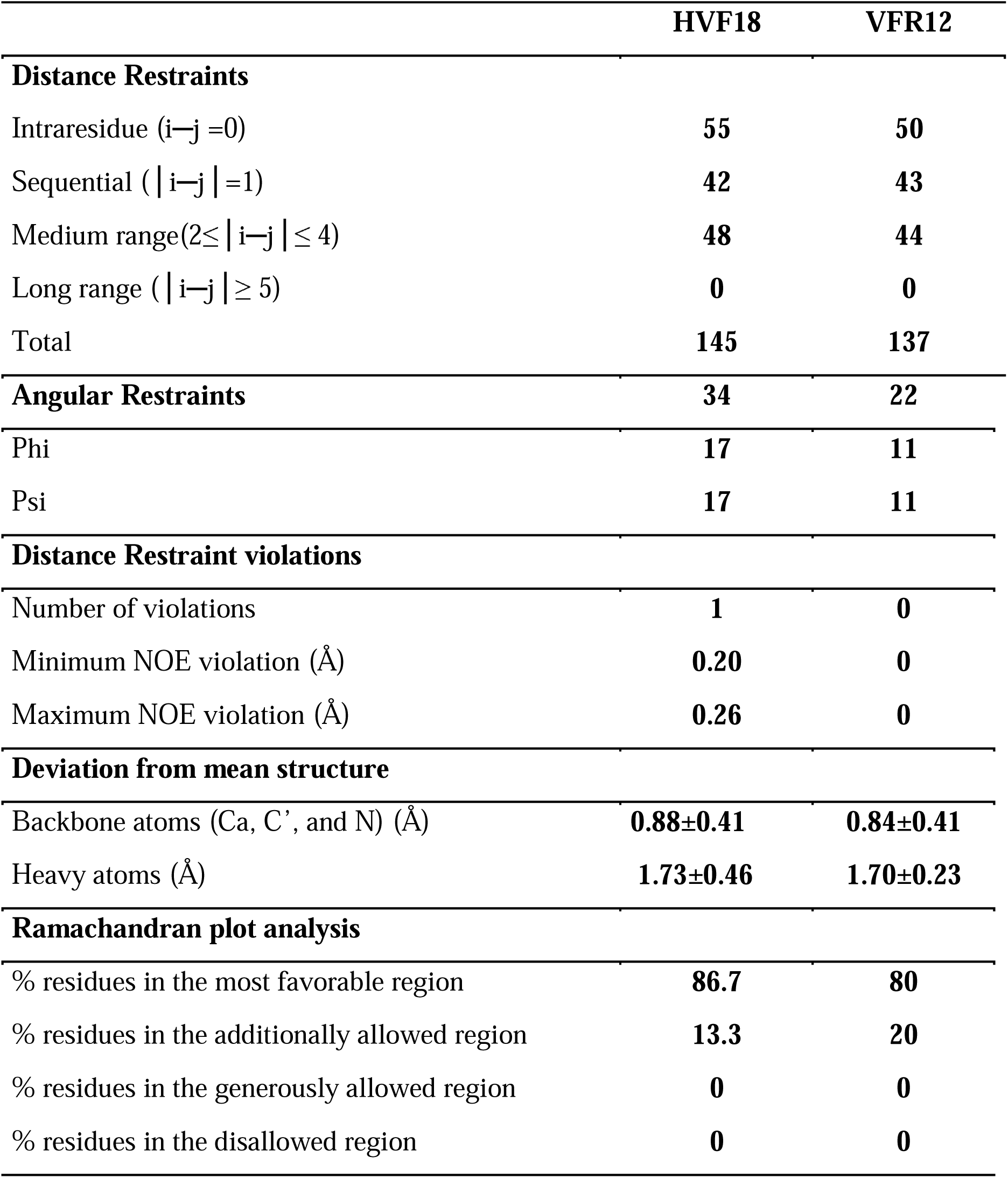
Summary of the structural statistics for the 20 lowest energy structures of HVF18 and VFR12 in LPS micelles

Likewise, the LPS-bound structure of VFR12 was determined using the tr-NOEs distance restraints (Fig. S5 and Table 2). The structure is well-defined with a backbone and heavy atom RMSD of 0.84 ± 0.34 and 1.70 ± 0.23 Å, respectively (Fig S5C). As in HVF18, the N-terminal residues (F2-L4) are extended with a β-turn (K5-I8), followed by an extremely short C-terminus helical turn I8-V11 (Fig S5B). The formation of the helical segment is confirmed by the diagnostic medium-range HN/HN (*i* to *i*+2), Cα/NH (*i* to *i*+2, *i*+3 and *i*+4) contacts for the C-terminal residues (Fig S5A). Consequently, the overall structure of LPS bound VFR12 is linear and the amphipathic distribution of side chains is disturbed. Although a hydrophilic surface including R3/K5/K6/Q9/K10 is formed in VFR12, the hydrophobic patch is small, involving only side-chains of F2/W7/V11 (Fig S5E-F).

### Simulated assembly of TCPs with lipid A aggregates

Next, tr-NOESY derived HVF18 and VFR12, as well as modeled GKY25, were allowed to bind (at a 1:2 ratio of peptide:lipid) to lipid A aggregates comprising 60 molecules during coarse-grained (CG) molecular dynamics (MD) simulations (Fig. S6). Lipid A was chosen for computational efficiency in the simulations, as it is the core effector component of LPS [18]. Divalent Ca^2+^ ions included in the simulations served to cross-link phosphate moieties between lipid A molecules, and are known to be essential for the stability of the resultant lamellar phase membranes and other biologically relevant phases [28, 29]. At t=250 ns, large clusters of peptides were formed on the surface of the lipid aggregates. Eventually, all of these peptide clusters dispersed into the aggregate over the course of each 10 μs simulation (Fig. S6A). The positively charged N-terminal residues were observed to compete with the Ca^2+^ ions that cross-linked between the phosphates of lipid A, resulting in more loosely connected headgroups (Fig S6B). This “breaking up” of the headgroup-crosslinking interactions occurred concurrently with the dispersion of the peptide clusters into the lipid aggregates. The dispersion of VFR12 was more rapid than that of HVF18 and GKY25, both of which formed more stable clusters that did not fully disperse within the 10 μs of simulation. All three peptides, were able to interact with multiple lipid molecules simultaneously (Fig S6B). Indeed, both GKY25 and HVF18 were most often found to interact simultaneously with five separate lipid A moieties. In contrast, the shorter VFR12 appeared to be less able to interact with multiple lipids, preferentially binding to four separate molecules corresponding to the reduced ability of VFR12 to neutralize LPS (Fig. S1D). The hydrophilic and positively charged residues at the N-terminus, K2, T7, H8* R11*, K13* and K14* of GKY25 (* also in HVF18) were found to be in contact with only the head group particles of lipid A (Fig S7A). The aromatic residues, Y3, F5, Y6, and F10*, are interfacial residues, binding both the head and tail parts of the lipid, while the C-terminal F22* was more often in contact with the tails. The hydrophilic and negatively charged residues of the C-terminal helix were solvent exposed, and had a propensity to interact with the Ca^2+^ ions after they had been displaced from their sites crosslinking lipid phosphates. Furthermore, high resolution all-atom (AA) molecular dynamics (MD) simulations of VFR12 and HVF18 served to validate the coarse-grained (CG) simulations (Fig 5A). These additional simulations revealed a similar mode of binding to lipid A moieties as observed in the CG simulations, with the N-terminal positively charged residues serving to make contacts with headgroups, and the hydrophobic residues of the C-terminal helix making contacts with the lipid interfacial region and tails atoms (Fig. 5B). Taken together, the combined CG and AA MD simulations correlate with the structural and *in vitro* effect studies, providing a detailed description of TCP–LPS interactions as well as regions directly contacting LPS that is reflected in the LPS neutralizing capabilities of the different TCPs.

**FIGURE 5.**
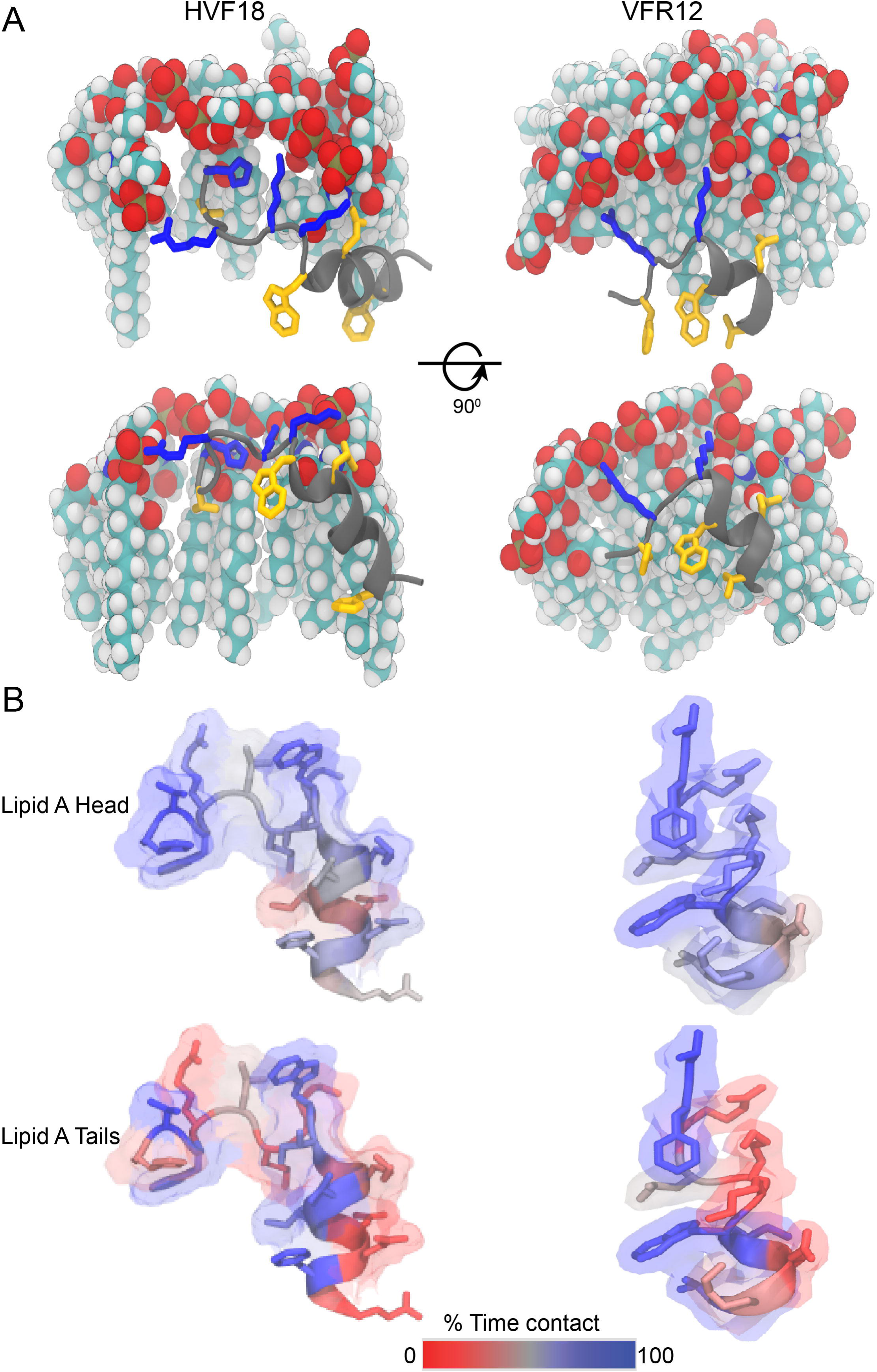
Molecular dynamics (MD) simulations of thrombin C-terminal peptides with lipid A aggregates. (A) All-atom (AA) MD simulations of HVF18 and VFR12 showing side chains of hydrophilic (blue) and hydrophobic (yellow) residues in contact with the headgroup and lipid A acyl tail. (B) HVF18 and VFR12 NMR structures highlighting the side chain residue contact differences with the LPS head group and acyl chain obtained from coarse grained (CG) molecular dynamics simulations.

### *In silico* analysis and microscale thermophoresis studies of LPS and TCP interaction with CD14

CD14 is a well-known pattern recognition receptor that plays a prominent role in sensitizing cells to LPS, and transferring it to the TLR4 signaling complex [30]. Considering the ability of TCPs to bind LPS and block TLR4 dimerization at cell surfaces [6, 24], *in silico* analyses were performed to investigate possible TCP-CD14 interactions. Mutational studies suggest that charged surfaces at the amino-terminus of CD 14 may be used to “capture” LPS, and the hydrophilic “rim” in particular to be of importance for LPS binding and cell activation [31]. In the absence of a ligand-bound structure for CD14, unbiased all-atom MD simulations were performed with the previously reported crystal structure of human CD 14 [32], to which a nearby lipid A molecule spontaneously bound to the suggested binding cavity (Fig. 6A). Subsequently, the energetic properties of binding a lipid A molecule to the hydrophobic pocket were investigated (Fig. S8) using biased all-atom MD simulations, based on an umbrella sampling (US) methodology [33]. The resultant potential of mean force (PMF) provides a measure of the lipid binding free-energy as a function of distance from the CD14 binding cavity [34]. The calculated PMF displays a smooth rise in free-energy upon extraction of lipid A from the CD14 binding cavity towards the bulk solvent phase, plateauing ~2 nm from the cavity (Fig. S8A). The absolute free-energy change between the CD14-bound and fully solvent-dissolved states was ~200 kJ mol^-1^ (Fig. S8B). This is similar in magnitude to the previously calculated binding free-energy change of lipid A to the structurally characterized LPS binding cavity of MD-2, the co-receptor of TLR4 [35], hence confirming that the proposed N-terminal location of CD14 is indeed a true binding site for LPS molecules.

**FIGURE 6.**
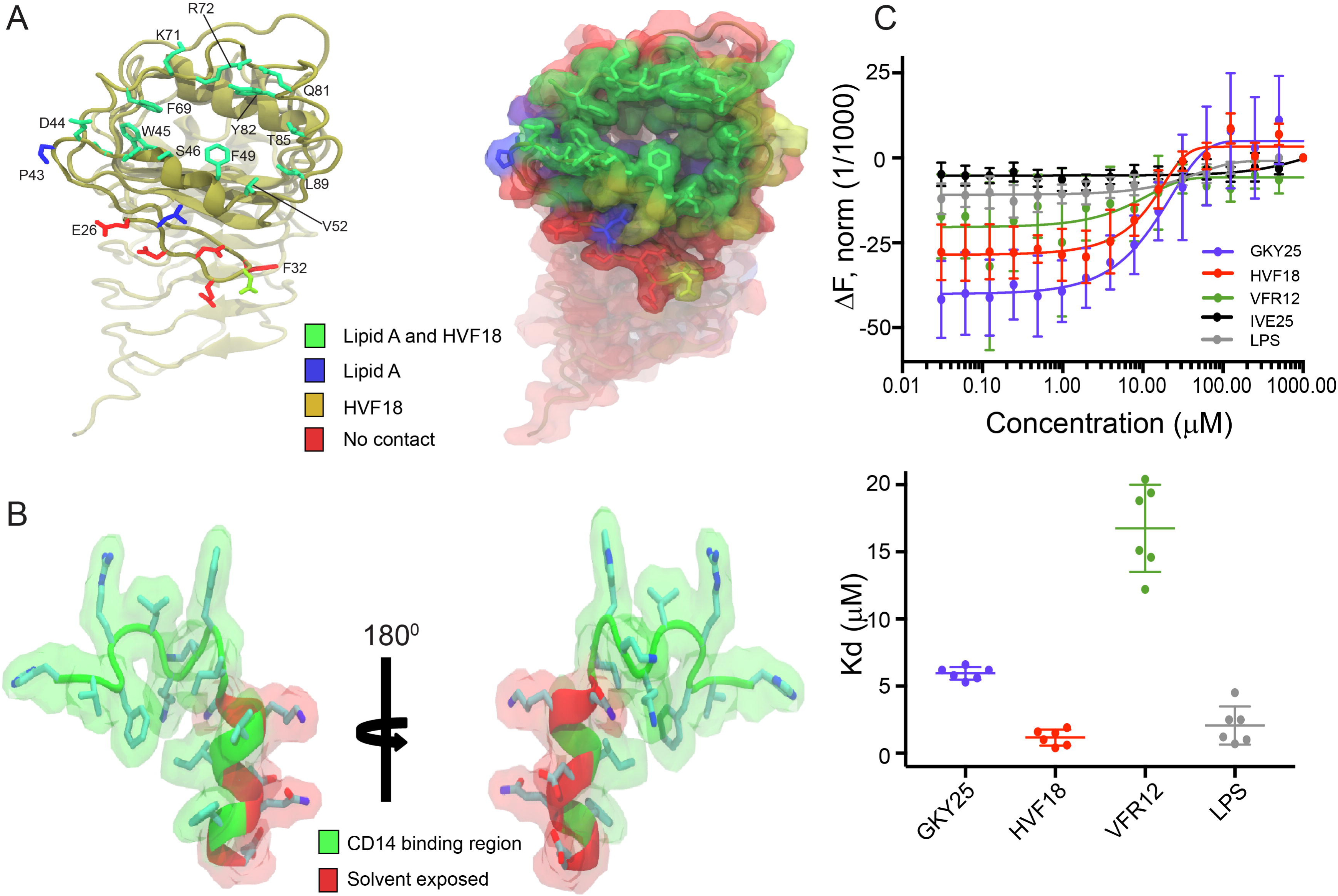
Thrombin C-terminal peptides interaction and binding to human CD14 ligand binding pocket. (A) Ribbon representation of CD 14 N-terminal hydrophobic pocket rim highlighting side-chains of residues contacting LPS (blue), HVF18 (yellow) and both LPS and HVF18 (green) in an all-atom molecular dynamics simulation model. (B) Ribbon representation of HVF18 indicating residues in contact with CD14 hydrophobic pocket (green) and solvent exposed (red). (C) Microscale thermophoresis assay quantifying CD14 interaction with TCPs and LPS. The peptide IVE25 was used as negative control. MST analysis were performed in concentration range 1000 – 0.03 μM of HVF_18_, GKY_25_IVE_25_ and 500 - 0.03 of LPS. (B) Kd constants of HVF_18_=1.2 μM, GKY25= 6.0 μM, VFR12= 16.8 μM and LPS=2.1 μM were determined. The mean values of six measurements ± their standard deviations are shown.

Docking of tr-NOESY derived HVF18 to human CD14 using the ClusPro server [36] indicated the peptide to be bound to the amino terminal hydrophobic pocket of CD14 in >90% of the docked models retrieved from the server (Fig. 6). In majority of these structures, the N-terminal tail of HVF18 was buried deeply into the hydrophobic pocket of CD14, making clear contacts with rim residues W45, F49, V52, F69 and Y82 that bind LPS tails during simulations (Fig 6A). In contrast, the C-terminal helix of the peptide bound to variety of hydrophilic rim residues, such as K71, R72, and Q81, as well as to rim residues specific for human CD14, such as the D44, S46, T85, and L89 (Fig 6A). In all the docked poses, HVF18 would contact CD14 in a common orientation with the C-terminal residues of Q10, K11, D14, Q15, and E18 exposed to the solvent. In contrast, VFR12 had a less specific mode of interaction, binding to CD14 in a variety of orientations. In a few cases, VFR12 would lie across the entrance to the hydrophobic pocket, making contact with all the rim residues of CD14. A GKY25-bound state was inferred by modeling the N-terminal residues as a random coil, using HVF18 as an initial template. The highest quality models of the GKY25-CD14 complex, i.e., those with the lowest values of the objective energy function, would either loop the N-terminal residues into and out of the pocket (Fig S7B), or lay at the N-terminus across the entrance to the CD14 pocket, completely blocking entry.

Likewise, microscale thermophoresis (MST), a highly sensitive technique probing interactions between components in solution, validated the *in silico* simulations presented above. As can be seen in Figure 6D, fluorescence-labeled CD14 binds *E. coli* LPS, with a K_d_ of 2.1 ± 1.4 μM, corresponding with previous reports [3]. Interestingly, both HVF18 and GKY25 showed a significant affinity to CD14 (K_d_ of 1.2 ± 0.6 and 6.0 ± 0.5 μM, respectively), contrasting the lower interaction observed for VFR12 (K_d_ of 16.8 ± 3.2 μM). Overall, these results indicate that the endogenous HVF18 and prototypic GKY25 efficiently block LPS-induced downstream signaling through direct LPS sequestration as well as high affinity antagonist binding to the amino terminal hydrophobic binding pocket of CD14, blocking LPS CD14 interactions.

## Discussion

Today, there is an increasing interest in developing HDPs as immune modulators [11, 37, 38], and natural sequences such as the TCPs studied here are of interest for use in human therapy. In this perspective, it is important to define peptide actions at the molecular level for rational design and development of new peptide-based anti-infectives [39-42]. The HVF18-LPS structure determined here comprises a C-terminal helix and extended N-terminus with a β-turn (Fig. 4) that favors a curved backbone fold, imparting a C-shaped amphipathic arrangement of the hydrophobic core lined with two hydrophilic clusters. Moreover, the lack of tr-NOEs in GKY25-LPS and FYT21-LPS complexes, implies that the N-terminal sequence (GKYGFYT) comprises the high affinity LPS-binding segment of TCPs (Fig. 1 and S3), in agreement with the effect of hydrophobicity in augmenting LPS neutralizing effects [43] as well as antimicrobial activity [44, 45]. Our MD simulations provided detailed visualization into TCPs interaction with LPS and CD14 (Fig. 5). The combined CG and AA molecular dynamics simulations identified key residues critical for the anti-inflammatory activity of TCPs (Fig 6). In contrast to HVF18 and GKY25, VFR12 lacks the N-terminal hydrophobic residues as well as the C-terminal hydrophilic residues that display increased contacts with the LPS head and acyl chain. Consequently, VFR12, though capable of still binding to LPS (via R3, K5, K6) comprises a relatively small hydrophobic patch, making it inefficient in penetrating the hydrophobic lipid phase of the aggregates in solution as well as interfering with LPS binding to the pocket of CD14 (Fig S5).

Over the years, tr-NOESY has been employed to obtain three-dimensional structures of LPS-interacting peptides such as lactoferrin [46], LBP [47], the magainin analogues MSI-78 and MSI-594 [48], and pardaxin [49], providing information on their mechanism of LPS-neutralization and bacterial outer membrane permeabilization [46-50]. The wide diversity of backbone folds in these structures highlights a certain degree of variability among LPS-interacting structures [51]. For example, pardaxin or MSI-594 forms helix-loop-helix structures in LPS, while lactoferrin derived LF-11, LBP derived LBP-14 or a *de novo* designed YW12 [39] form β-hairpin or β-boomerang structures, respectively. The heparin cofactor II-derived peptide KYE28 displays an N-terminal short helical region followed by a loop, two turns, and an extended C-terminus when bound to LPS [52]. Nevertheless, a feature of these determined structures is that they display partly well-defined and flexible structures in complex with LPS, characterized by an amphipathic fold [53]. In this perspective, it is of interest that TCPs share many of these overall characteristics, although they form a unique C-shaped structure, reminiscent of the T-shaped fold of LF11 (Fig. S9). Interestingly, the TCP-region RLKKWI contains the cationic and aromatic amino acid residues proposed as critical for lipid A interactions [54], and present in several LPS-binding proteins [55]. From an evolutionary perspective, it is notable that these residues show a high degree of conservation among various thrombin species. It is also notable that these residues are part of the region which binds to the LPS-binding pocket of CD14, whereas the less conserved residues in the C-terminal end are outside this pocket (Fig. S10A).

From a biological perspective, we have previously shown that TCPs such as GKY25 and FYT21 interact and bind directly to macrophages/monocytes *in vivo,* leading to inhibition of TLR4 dimerization and subsequent abrogation of pro-inflammatory cytokine production [6, 24]. Although these data hinted at the possibility that TCPs may interact with targets other than LPS, the mechanistic basis for the anti-inflammatory effects of such peptides remained unclear. Using *in silico* docking and MST analysis, we here show that TCPs bind to the amino terminal hydrophobic LPS binding pocket of human CD14, the primary receptor of LPS with high affinity (Fig. 6). This prevents LPS transfer to TLR4 and consequently reduces macrophage activation. Secondly, the regions identified to interact with LPS overlap with the residues involved in the interaction with CD14, suggesting that TCPs can interfere with LPS-CD14 interactions, hence abrogating downstream transfer of LPS to the MD-2 complex and other effectors. It is also notable that HVF18 [6, 23], binds CD14 with an affinity similar to that obtained for LPS. In this context, it is therefore interesting that multiple variants of TCPs are generated by human and bacterial proteases, several of which are found in human wound fluids (Fig. S10B) [23]. This observation implies that shorter TCP fragments, related to VFR12, may be exclusively binding to LPS and bacteria, whereas longer fragments like the HVF18 or FYT21 have the capacity also to neutralize LPS and inhibit TLR4 signaling pathways. Furthermore, the fact that the major TCP fragment generated by human neutrophil elastase, HVF18, shows a higher affinity for CD14 than the longer GKY25, whereas the reciprocal applies to their interactions with LPS, illustrates that protease-mediated generation of multiple TCPs may “tune” LPS-responses during wounding and infection. It is also interesting that the affinities between TCPs and CD14 were in the |jM range, similar to the K_d_ of LPS to CD14. Hypothetically, this could enable TCPs to down-modulate excessive responses to bacteria and LPS responses, without completely abrogating such responses, as they also have a physiologically relevant function in infection clearance. Indeed, such phenomena have been observed in experimental models of bacterial sepsis and endotoxic shock, where treatment with TCPs reduced cytokine responses and animal mortality [4, 7]. It has been increasingly appreciated that a multitude of weak, or transient, biological interactions (K_d_ > μM), either working alone or in concert, occur frequently throughout biological systems [56, 57]. The here disclosed TCP family, with its multiple and relatively weak affinities, all in the μM range, may represent an elegant example of such an endogenous system, with an ultimate function of modulating host responses to infection. Sharing many characteristics with ‘transient drugs’, defined by their multivalency, multiple targets, high-off-rates and K_d_ rates at μM levels [56], TCPs could therefore be of interest in the development of novel antiinflammatory therapies.

## Materials and Methods

### Peptides

The peptides GKY25 (GKYGFYTHVFRLKKWIQKVIDQFGE), FYT21 (FYTHVFRLKKWIQKVIDQFGE), HVF18 (HVFRLKKWIQKVIDQFGE), and VFR12 (VFRLKKWIQKVI), were synthesized by Biopeptide (San Diego, CA). We confirmed the purity (over 95%) using mass spectral analysis (MALDI-TOF Voyager, USA).

### Digestion of α-thrombin and Western blotting

Human α-thrombin (4 μg) (Innovative Research, Inc, USA) was incubated with human neutrophil elastase (HNE) (Calbiochem, USA) at an enzyme-substrate ratio of 1:30 (w/w) in 10 mM Tris, pH 7.4 (volume of 10 μl) at 37 °C for 60 and 180 minutes. After incubation, the enzyme was inactivated by heating the samples at 95 °C for 3 minutes. After addition of 7 μl of Novex sample buffer and heating at 95 °C for 10 minutes, samples were analyzed using 10-20% Novex Tricine pre-cast SDS-PAGE gel (Invitrogen, USA) run at 90 V, for 2 hours. Immediately following electrophoresis, western blotting was performed at 25V for 90 minutes on ice as mentioned earlier with few modifications [7]. After the transfer, PVDF membrane (Bio-Rad, USA) was blocked with 5% milk for an hour, followed by overnight incubation with rabbit polyclonal antibodies recognizing the thrombin C-terminus peptide sequence (VFRLKKWIQKVIDQFGE) (Innovagen, Lund, Sweden) in 1:5000 dilution at 4 °C. Post washing (3 x 10 minutes) the membrane was incubated with HRP-conjugated secondary antibodies (Dako, Denmark) in 1:10000 dilutions for an hour. The membranes were again washed 3 times for 10 minutes and thrombin C-terminal fragments were detected by SuperSignal West Pico Chemiluminescent Substrate (ThermoFisher Scientific, USA), according to manufacturer recommendations and thereafter imaged using a ChemiDoc MP System (Bio-Rad, USA).

### Fluorescence spectroscopy

Fluorescence experiments were carried out with a Cary Eclipse fluorescence spectrophotometer (Varian Inc., Australia) equipped with dual monochromators using a 0.1 cm path length cuvette and a slit width of 5 nm. All fluorescence spectra were recorded in 10 mM Tris buffer, pH 7.4. Intrinsic tryptophan fluorescence spectra of free peptide were recorded using 5 μM peptide. Interaction between peptide and LPS was studied by titrating the peptide with increasing lipid concentration up to 10 μM. The intrinsic tryptophan was excited at a wavelength of 280 nm and the emission monitored between 300-400 nm. The changes in emission maxima were further used to determine the binding affinity of the peptides with the LPS using Graph Pad Prism v7.02 (GraphPad Software, Inc. USA), assuming a single binding site. Fluorescence quenching of the tryptophan residue was performed by adding increasing concentrations of quencher (acrylamide) (stock 5 μM) into 10 mM Tris buffer, pH 7.4, containing either unbound (5 μM) peptide or in the bound state containing saturating concentration of LPS (10 μM). Quenching of fluorescence intensity at emission maxima for corresponding free and lipid-bound forms was analyzed using the Stern-Volmer equation F_o_/F= 1+KSV(Q), where F_o_ and F are the fluorescence intensity at emission maxima in the absence and presence of quencher, respectively, K_SV_ is the Stern-Volmer quenching constant, and [Q] is the molar concentration of quencher [50].

### Ellipsometry

Peptide adsorption was studied *in situ* by null ellipsometry [58, 59], using an Optrel Multiskop (Optrel, Kleinmachnow, Germany) equipped with a 100 mW Nd:YAG laser (JDS Uniphase, Milpitas, USA). All measurements were carried out at 532 nm and an angle of incidence of 67.66° in a 5 ml cuvette with stirring (300 rpm). In brief, by monitoring the change in the state of polarization of light reflected at a surface in the absence and presence of an adsorbed layer, the mean refractive index (n) and layer thickness (d) of the adsorbed layer can be obtained. From the thickness and refractive index, the adsorbed amount (Γ) was calculated according to:

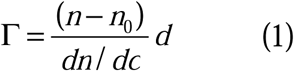

where dn/dc (0.154 cm^3^/g) is the refractive index increment and n_0_ is the refractive index of the bulk solution. Corrections were routinely done for changes in bulk refractive index caused by changes in temperature and excess electrolyte concentration.

LPS-coated surfaces were obtained by adsorbing *E. coli* LPS to methylated silica surfaces (surface potential -40 mV, and contact angle 90°[60]) from 5 mg/ml LPS stock solution in water at a concentration of 0.4 mg/ml in 10 mM Tris, pH 7.4. This LPS concentration corresponds to saturation in the LPS adsorption isotherm under these conditions. Non-adsorbed LPS was removed by rinsing with Tris buffer at 5 ml/min for a period of 30 minutes, allowing the buffer to stabilize for 20 minutes. Peptide addition was subsequently performed to 0.01, 0.1, 0.5, and 1 μM, and adsorption monitored for at least one hour after each addition. All measurements were performed in at least duplicate at 25 °C.

### Limulus amebocyte (LAL) assay

The ability of peptides to sequester LPS was assayed using a commercially available LAL chromogenic kit (QCL100 Cambrex, Lonza, USA), according to the manufacturer’s protocol. LPS activates a proenzyme in the *Limulus* amebocyte lysate (LAL), which in turn catalytically splits a colorless substrate to the colored product para-nitroanilide (pNA), which is measured spectrophotometrically at 410 nm. Prior to running the assay, the peptides were dissolved at 200μM in pyrogen-free water, supplied with the kit.

### NF-κB activation assay

THP1-XBlue-CD14 reporter cells (1x10^6^ cells/mL) (InvivoGen, San Diego, USA) were seeded in phenol red RPMI (180,000 cells/well), supplemented with 10% (v/v) heat-inactivated FBS and 1% (v/v) Antibiotic-Antimycotic solution (AAS), and allowed to adhere before they were stimulated with 100 ng/mL *Escherichia coli* (0111: B4) LPS (Sigma, USA) and with peptides at the indicated concentrations. The NF-κB activation was determined after 20 h of incubation according to the manufacturer’s instructions (InvivoGen, San Diego, USA). Briefly, activation of NF-κB leads to the secretion of embryonic alkaline phosphatase (SEAP) into the cell supernatant, where it is measured by mixing the supernatant with a SEAP detection reagent (Quanti-Blue™, InvivoGen), followed by absorbance measurement at 600 nm. Data shown are mean values obtained from triplicate measurements.

### CD spectroscopy

The secondary structure of unbound and LPS bound peptides was analyzed by circular dichroism (CD) spectroscopy using a Chirascan Circular Dichroism spectrometer, (Applied Photophysics, U.K.) with a cuvette of path length of 0.05 cm (Hellma, GmbH & Co, Germany). Spectral data were collected at a step size of 0.5 nm, time constant of 1s and peptide concentration of 20 μM in 10 mM Tris, pH 7.4. Structural changes upon addition of *E. coli* 0111:B4 LPS (100 μM), was also recorded in the same buffer. The molecular weight of LPS was considered to be 10 kDa, according to a previous report [61]. All spectra were recorded at 25°C from 190 to 260 nm, using a bandwidth of 1-nm, and averaged over three scans. Baseline scans were obtained using the same parameters for buffer or LPS and subtracted from the respective data scans with peptides. The final corrected averaged spectra were expressed in mean residue ellipticity.

### NMR Spectroscopy

All NMR experiments were performed on a Bruker DRX 600 MHz spectrometer (Bruker, Germany), equipped with cryo-probe and pulse field gradients. NMR data processing and analysis was carried out using Topspin software (Bruker) and the SPARKY NMR assignment program (https://www.cgl.ucsf.edu/home/sparky). Twodimensional ^1^H-^1^H TOCSY and ^1^H-^1^H NOESY spectra of GKY25, FYT21, HVF18, and VFR12 were acquired at 298K at a peptide concentration of 0.5 mM in aqueous solutions containing 10% D_2_O, pH 4.5 at a mixing time of 80 and 200 ms respectively. 4,4-dimethyl-4-silapentane-5-sulfonate sodium salt (DSS) was used as an internal standard (0.0 ppm) in all NMR experiments. The peptide-LPS interaction was monitored by a series of one dimensional ^1^H-NMR spectra titrated with various concentrations of LPS until line broadening was observed. Two-dimensional tr-NOESY experiments for VFR12 and HVF18 were acquired in the presence of 20 μM and 30 μM LPS with mixing times of 100, 150 and 200 ms. The peptide-LPS complex was stable over the period of data collection and generated a large number of tr-NOE cross-peaks. All two-dimensional tr-NOESY experiments were performed with 400 increments in t1 and 2048 data points in t2 using WATERGATE [62] procedure for water suppression and States-TPPI [63] for quadrature detection in the t1 dimension. The spectral width was 12 ppm in both dimensions. After 16 dummy scans, 80 scans were recorded per t1 increment. After zero filling in t1, 4K (t2) × 1K (t1) data matrices were obtained.

### NMR-derived structure calculations

The LPS-bound three-dimensional structures of VFR12 and HVF18 were calculated using the CYANA program (version 2.1) [64] from tr-NOESY (mixing time 150 ms) derived distance constraints. NOE intensities were qualitatively assigned to strong, medium, and weak cross-peak intensities with upper bound distance limits of 3.0, 4.0, and 5.5 Å, respectively. The lower bounds for all distance restraints were fixed to 2.0 Å. The backbone dihedral angle (Phi) was varied between -20° and -180° to reduce conformational sampling [39, 65]. No hydrogen bond constraints were used for structure calculation. Several rounds of structure calculations were performed and depending on the NOE violations, the angle and distance constraints were refined. Finally, out of 100 generated structures, the 20 lowest energy conformers with low RMSD were selected for the NMR ensemble. Structures were visualized using the MOLMOL and PyMOL software. The quality of the determined structure was validated using PROCHECK-NMR [66].

### Peptide assembly simulations to LPS aggregates

All molecular dynamics (MD) simulations were performed using the GROMACS simulation package [67, 68]. All-atom (AA) MD simulations of VFR12 and HVF18 were performed using the CHARMM27 force field including additional CMAP potentials [69, 70]. Electrostatic interactions were treated using the smooth Particle Mesh Ewald (PME) method [71], with a cutoff for short-range interactions of 1.2 nm. The van der Waals interactions were switched to zero between 1.0 nm and 1.2 nm. The system temperature was maintained at a constant temperature of 313 K using the velocity-rescale thermostat and a coupling constant of 1 ps[72]. A pressure of 1-bar was maintained isotropically using the Parrinello-Rahman barostat and a coupling constant of 5 ps.

To ensure that the conformational space accurately reproduced the tr-NOESY derived structures, the NOEs were modeled in the simulations as flat-bottomed distance restraint potentials with a force constant of 1000 kJ mol^-1^ nm^-2^. The time constant for the distance restraints running average was 10 ns. Angle restraints of 100 kJ mol^-1^ rad^-2^ were also applied to maintain the experimentally-derived backbone conformation. A lipid A aggregate was created by manually positioning ten lipid A molecules in solution using VMD [73] and allowing them to self-assemble over a timescale of 100 ns. The lipid parameters were used as described previously [74, 75]. The peptides were positioned ~2.0 nm from the aggregate surface. Energy minimization, using the steepest descent algorithm, was performed for 1000 steps. Position restraints, with a force constant of 1000 kJ mol^-1^ nm^-2^, were placed on the Cα particles of the peptides for 5 ns. Production simulations were performed for 250 ns.

CG MD simulations were performed using the MARTINI 2.2 force field [76]. DSSP was used to determine the secondary structures of the tr-NOESY derived VFR12 and HVF18 structures. The DSSP secondary structure categories served as constraints in the preparation of MARTINI protein parameters [77, 78]. The additional seven residues at the N-terminus of GKY25, with respect to HVF18, were modeled as a random coil. The large lipid A aggregate was created by allowing 60 randomly placed lipid molecules to assemble over the course of a 1 μs simulation. The final frame of this simulation was used as the starting point for the creation of three further systems in which 30 GKY25, HVF18 or VFR12 peptides, respectively, were scattered randomly in the simulation cell, all at least 2 nm from the aggregate. The GKY25, HVF18 and VFR12 CG production simulations were each run for 10 μs. A simulation of the aggregate alone, with no peptides added, was also extended to 11 μs, and this served as negative control. All CG simulations were maintained at a constant temperature of 313 K using velocity rescale thermostat [72]. A system pressure of 1 bar was maintained isotropically using the Parrinello-Rahman barostat [79, 80] with a coupling constant of 20 ps. Electrostatic interactions were treated using the reaction-field method [81] with a dielectric constant of 15 within a 1.1 nm cutoff, and zero beyond the cutoff. The van der Waals interactions were shifted to zero at 1.1 nm.

### Peptide and LPS interference with CD14 interactions

The tr-NOESY derived structures of VFR12 and HVF18 were docked to human CD14 using the ClusPro server [82]. The structure of CD14 with GKY25 bound was modeled using Modeller v9.15 [83] with the HVF18-CD14 complex model as a template via the loopmodel function with the MD refinement set to “refine.very_slow”. A total of 100 models of the GKY25-CD14 complex were generated. The structure with the lowest value of Modeller’s objective function was selected for visualization in Figure 6. In order to validate the GKY25-CD14 model, GKY25 was also modeled in isolation using tr-NOESY derived structure of HVF18 as a template. One hundred models were generated, and the model with the lowest objective function was selected for further docking studies. The modeled GKY25 structure was docked to human CD14 using the ClusPro server.

Lipid A was assembled in an unbiased mode to CD14 via molecular simulations. Initially, a single lipid A molecule was placed ~2.0 nm from the binding pocket (measured from the center of mass of the terminal acyl carbons of lipid A to the center of the cavity). Rapid spontaneous adsorption and insertion of lipid A into the CD14 N-terminal binding cavity was observed (within 10 ns). The simulation was subsequently run for over 1 μs and the lipid molecule remained stably bound within the N-terminal pocket.

For free-energy calculations, the lipid A bound CD14 structure was extracted after 1 μs of simulation. The protein and ligand were manually positioned so that pulling the lipid A molecule along the unit cell z-axis would correspond to the axis of the cavity mouth. The system was then re-solvated and minimized before a 4 ns steered MD simulation was performed, pulling the lipid A molecule 4 nm from the protein using a pulling force of 1000 kJ mol^-1^ nm^-2^. Forty windows were extracted at 0.1 nm intervals from this trajectory for subsequent umbrella sampling (US). During US, each window was sampled for 50 ns. The lipid A sugar backbone and phosphate groups served as the reference point of the ligand, and the backbone atoms of residues 8 to 12 and 31 to 35 corresponding to the first two LRRs at the cavity entrance, were used as the reference point for CD14.

### Microscale thermophoresis (MST)

MST analysis was performed using a NanoTemper Monolith NT.115 apparatus (Nano Temper Technologies, Germany). A Monolith NT Protein labeling kit RED – NHS (Nano Temper Technologies, Germany) was used to label 2μM of sCD14 (PeproTech, USA) according to the manufacturer’s protocol. A constant amount of 0.5 of sCD14 was incubated for 30 minutes at room temperature with increasing concentrations of VFR12, HVF18, GKY25 (1000 - 0.03 μM) or LPS (500 – 0.03 μM) in Tris buffer (10 mM, pH 7.4). Next, 10 μl of the sample was loaded into standard glass capillaries (Monolith NT Capillaries, Nano Temper Technologies), and the MST analysis was performed (settings for the light-emitting diode and infrared laser were 80%). Labeled IVE25 was added to sCD14 under the same conditions as above and used as negative control.

## DATA DEPOSITION

The atomic coordinates and NMR constraints of LPS bound HVF18 and VFR12, energy minimized conformers, are deposited to Biological Magnetic Resonance data Bank (BMRB) with the accession codes 21081 and 21080 respectively.

## ACKNOWLEDGEMENTS

This research was supported by the Lee Kong Chian School of Medicine, Nanyang Technological University Start-Up Grant, the Lee Kong Chian School of Medicine Postdoctoral Fellowship (RS), Nanyang Technological University Singapore Ministry of Education under its Singapore Ministry of Education Academic Research Fund Tier 1 (2015-T1-001-082), the Swedish Research Council (project 2012-1883), the Knut and Alice Wallenberg Foundation, and the Swedish Research Council (projects Swedish Research Council (2016-05157) (MM) and 2012-1883 (AS)), as well as by the LEO Foundation Center for Cutaneous Drug Delivery (2016-11-01) (MM). Ann-Charlotte Strömdahl is gratefully acknowledged for skillful technical support.

